# Principal component of explained variance: an efficient and optimal data dimension reduction framework for association studies

**DOI:** 10.1101/036566

**Authors:** Maxime Turgeon, Karim Oualkacha, Antonio Ciampi, Golsa Dehghan, Brent W. Zanke, Andréa L. Benedet, Pedro Rosa-Neto, Celia MT. Greenwood, Aurélie Labbe, for the Alzheimer’s Disease Neuroimaging Initiative

## Abstract

The genomics era has led to an increase in the dimensionality of the data collected to investigate biological questions. In this context, dimension-reduction techniques can be used to summarize high-dimensional signals into low-dimensional ones, to further test for association with one or more covariates of interest. This paper revisits one such approach, previously known as Principal Component of Heritability and renamed here as *Principal Component of Explained Variance* (PCEV). As its name suggests, the PCEV seeks a linear combination of outcomes in an optimal manner, by maximising the proportion of variance explained by one or several covariates of interest. By construction, this method optimises power but limited by its computational complexity, it has unfortunately received little attention in the past. Here, we propose a general analytical PCEV framework that builds on the assets of the original method, i.e. conceptually simple and free of tuning parameters. Moreover, our framework extends the range of applications of the original procedure by providing a computationally simple strategy for high-dimensional outcomes, along with exact and asymptotic testing procedures that drastically reduce its computational cost. We investigate the merits of the PCEV using an extensive set of simulations. Furthermore, the use of the PCEV approach will be illustrated using three examples taken from the epigenetics and brain imaging areas.

## 1 Introduction

In the ‘omics era, a considerable amount of data is now routinely collected to investigate the relationships between a set of covariates of interest (*X*) and genetic, molecular, or clinical outcomes (*Y*). As such, a substantial proportion of the methodological research developed in the last decade has focused on the numerous statistical challenges and computational issues highlighted by the joint analysis of such high dimensional correlated data. If we imagine a spectrum of models indexed by dimensionality, at one end, a model which attempts to accommodate all variables at once may lead to results which are difficult to interpret. At the other extreme of this spectrum, univariate models which treat each variable separately while ignoring the others may miss subtle signals arising from complex biological interactions and correlations. An intermediate approach would therefore either identify *a priori* relevant groups of variables *Y* on which to perform analysis, or reduce the dimensionality of the problem by summarizing the data into meaningful components. A popular example of dimension reduction is principal components analysis (PCA), which seeks linear combinations of the original data that explain the maximum amount of variance.

In contrast to studies with high-dimensional *covariates*, here we focus on studies where the goal is to investigate association between a set of (possibly high dimensional) correlated *outcomes* and one or more covariates of interest. In this context, several methods integrating both data reduction and association analysis simultaneously have been developed. For example, in a variant of principal component regression (PCR), a PCA analysis is performed on the set of outcomes *Y*, producing a smaller number of components, which are then used in an association test with the covariates of interest *X*. Although being widely used in practice, this method has very poor power in cases where the outcomes showing the greatest variability, hence those captured by the first principal components, are not the ones associated with the covariates of interest. This is of particular concern in genetic studies, where outcomes strongly influenced by the environment could easily have much larger variability than outcomes influenced by a single nucleotide polymorphism (SNP). Therefore, PCA in this context is likely to find components where environmentally driven traits have the highest weights (since they are highly variable), whereas the weights associated with genetically controlled traits may be very small if they are less variable across indidivuals. If an association analysis is then performed between such components 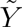 and genotype data *X*, it is likely that no association will be detected. Since the data reduction step in PCR is performed independently of the covariates, the low power in such situations is not surprising; as a consequence, other methods have been developed to jointly perform both data reduction and association analysis. For example, Partial Least Square (PLS) regression [1], Canonical Correlation Analysis (CCA) [2] or Linear Discriminant Analysis (LDA) [3] are widely used component-based methods that can be used to find, simultaneously, the “best” components describing two sets of variables (e.g. outcomes and covariates) by maximising the association between them. The optimisation criterion for association differs between these methods. For example, CCA is based on maximising correlations, whereas PLS optimises covariances. These types of methods are very powerful by construction, and have been extended in several ways to accommodate high dimensional variables using regularisation techniques or sparsity measures. However, such extensions heavily depend on tuning parameters, and formal testing procedures are currently lacking.

Although PLS and CCA have been gaining popularity in genomics, these methods were originally developed in other fields and mainly for prediction purposes. Almost twenty years ago, a closely related method implementing similar ideas to PLS and CCA was developed by Ott & Rabinowitz [4] specifically for a genetic context. Using family data and with the aim of performing linkage analysis between multiple correlated outcomes *Y* and SNP genotypes *X*, the idea was to find the best linear combination of outcomes (i.e. a component) maximising the heritability at the SNP tested. This method, termed Principal Component of Heritability (PCH), has unfortunately received little attention although it was later extended [5, 6, 7] to the more general setting of population-based studies and high-dimensional outcomes based on regularisation or sparsity techniques. Since heritability is simply defined in PCH as the proportion of variance in the outcomes *Y* explained by the SNP covariate, it can be seen as a method belonging to the family of PLS and CCA based on another criterion, i.e. the proportion of variance in the outcome variables explained by the covariates. Since we feel that, in the context of population-based studies, the term “heritability” is very misleading, and since the concept undelying PCH can be used also in a general setting outside the genetic field, we have renamed this approach Principal Component of Explained Variance (PCEV) We note that this method is also closely related to dual-scaling [8], introduced in the psychometrics literature.

In this paper, we present a completely new analytical framework based on the PCEV concept that copes smoothly with data of very high dimension at the outcome level, includes valid hypothesis tests, leads to interpretable results, and yet is computationally efficient. This approach is implemented in an R package, pcev. Specifically, we first show that unlike the competing, similar approaches discussed earlier, PCEV has mathematical properties that allow inclusion of very high dimensional outcomes without the need to rely on variable selection strategies that require choosing tuning parameters. Secondly, we show that PCEV can be extremely computationally efficient, unlike its competitors, by developing an exact testing procedure that does not rely on permutations. Therefore, this approach is entirely feasible for use in genome-wide and/or very high dimensional studies, such as we frequently encounter today.

We investigate the merits of the PCEV using an extensive set of simulations. As we want to focus our discussion on methods that do not require tuning parameters and those for which a testing framework is fully developped, we discarded from our comparison methods such as sparse PLS, regularized CCA, and the projection regression model proposed by Lin et al. [9] Furthermore, the use of the PCEV approach will be illustrated in three settings: i) using DNA methylation data derived from whole genome bisulfite sequencing (*Y*), we test if a region near the BLK gene on chromosome 8 is differentially methylated in B-cells compared to T-cells or monocytes; ii) using DNA methylation microarray data (*Y*), we perform a genome-wide gene-based association analysis of methylation with respect to cigarette smoking status; and iii) using brain imaging traits derived from [18F]Florbetapir PET scans (*Y*), we perform a genome-wide association study of amyloid-beta accumulation.

## 2 Method

The general methodological framework aims to simultaneously test a (possibly large) set of phenotypes or outcomes **Y**, against a set of covariates **X**. The method is evaluated through an extensive simulation study and then applied to three independent datasets.

### 2.1 General theoretical PCEV framework

We consider the following setting: let **Y** be a multivariate phenotype of dimension *p* (e.g. methylation values at *p* CpG sites, or brain imaging measured at *p* locations in the brain), let **X** be a *q*-dimensional vector of covariates of interest (e.g. smoking, cell type or SNPs) and let **C** be an *r*-dimensional vector of confounders (e.g. age or sex). We assume that the relationship between **Y** and **X** can be represented via a linear model

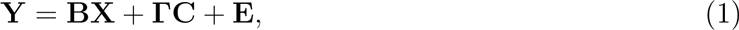

where **B** and **Γ** are *p × q* and *p × r* matrices of regression coefficients for the covariates of interest and confounders, respectively, and **E** ~ *N_p_* (0, **V**_R_) is a vector of residual errors. This model assumption allows us to decompose the total variance of **Y**, conditional on **C**, as follows:

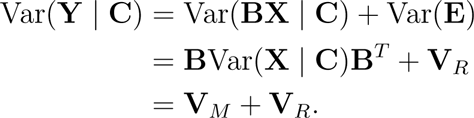

where 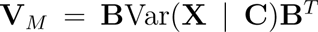 is the *model* variance component and **V**_R_ is the *residual* variance component. PCEV seeks a linear combination of outcomes, **w**^*T*^**Y**, which maximises the ratio *h*^2^(w) of variance being explained by the covariates **X**:

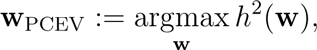

where

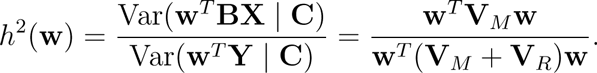

It has been shown that **w**_PCEV_ is the solution to the generalised eigenvector problem 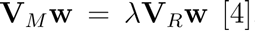 [4], and therefore standard linear algebraic results can be used to get a closed form for **w**_PCEV_. Although there are some similarities between PCA and PCEV since both methods seek a linear combination of outcomes optimising a given criterion, we recall that principal component analysis (PCA) reduces the dimension of **Y** by looking for a linear combination of its components with maximal variance, regardless of the covariates **X**. Furthermore, an important difference between PCA and PCEV is in the number of components one can extract. While the number of components to select in PCA is usually left to the user, and the maximal number of extracted components is bounded above by *p*, the maximal number of components that can be extracted for PCEV is bounded above by *q*, the number of covariates. Therefore, if we are only interested in one covariate, only one PCEV can be extracted; this follows from considering the rank of the matrix 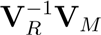.

### 2.2 PCEV with high-dimensional data

When the number of response variables, *p*, is larger than the sample size, *n*, a naïve implementation of PCEV will fail. To ensure the uniqueness of the solution to the maximisation process, the invertibility of the residual matrix **V***_R_* is required; therefore, an accurate estimation of the matrix **V***_R_* would require *n* > *p*. This limitation has led to the introduction of regularisation techniques for the estimation of **w**, similar in spirit to ridge regression [10] and Lasso [11]. We stress once again that these methods require parameters that are computationally expensive to compute. For this reason, we propose a novel alternative, namely a *block approach* to the estimation of PCEV. Assume we can partition **Y** into blocks (or clusters) such that the number of components in a given block is small enough, i.e. smaller than *n*. We can then perform PCEV and get a linear combination 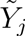 of the traits belonging to the *j*th block, for each block *j* = 1,…, *b*. We then obtain a new multivariate pseudo-phenotype 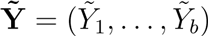, where each 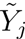 is of dimension one, on which we can again perform PCEV. Since the result is a linear combination of linear combinations, it is itself a linear combination of the original traits **Y**. Although one might think that this stepwise approach is an *ad-hoc* extension of the original PCEV approach, it has nonetheless a very appealing and relevant mathematical property, described in the following result:

**Theorem 1**. *Assume one can partition the outcomes* **Y** *into blocks in such a way that blocks are uncorrelated (i.e. outcomes lying in different blocks are uncorrelated). Then the linear combination (PCEV) obtained from the traditional approach and that obtained from the stepwise block approach described above are equal*.

The proof of this result is given in Appendix 1. Of course, if such a partition does not exist, there will generally be a difference between the two estimation procedures. Through a series of simulations and analyses of data sets, however, we will show that the discrepancy is small and comparable conclusions can be achieved even in the presence of some dependence between blocks.

### 2.3 Test of significance

The PCEV methodology was first presented in the context of family-based studies, and the lack of a proper significance testing framework may have hindered its adoption in population-based studies. Klei et al. [6] discussed such a framework, but their approach relies on computationally intensive sample splitting and resampling. In fact, here we are able to show that there is an analytic test of the null hypothesis *H*_0_: *h*^2^(**w**) = 0 that requires no resampling. This test is based on the largest eigenvalue λ of the matrix 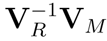, a test statistic used in multivariate analysis of variance [12]. When **X** consists of a single covariate, it is also known as the Wilks statistics; it can be shown that under the null hypothesis

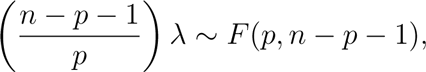

where *F*(*ν*_1_, *ν*_2_) is the Fisher-Snedecor distribution, with degrees of freedom *ν*_1_ and *ν*_2_ (see Appendix 2). The two matrices **V**_R_ and **V**_M_ are typically estimated during the classical PCEV process, which then allows us to compute the test statistic. When multiple covariates of interest are included in the model (e.g. multiple indicator variables for cell type composition, or multiple SNPs in a given genomic region), the statistic λ is better known as Roy’s largest root test statistic (also called Roy’s union-intersection test statistics) [13]. The asymptotic distribution of a suitable transformation of λ was derived by Johnstone [14]. More details are given in Appendix 3.

We note that the Wilks and Roy’s largest root test statistics both rely on the assumption of normality of the outcomes, and the former also requires *n* – *p* – 1 > 0 (corresponding to the second degree of freedom of the F test). If these assumptions are not satisfied, permutation tests provide an adequate control of the Type I error and can be used as an alternative.

### 2.4 Variable importance

The PCEV framework described above allows reduction of the multiple testing burden by performing only a single test for a set of outcomes. If the test is significant, it is of great interest to identify the set of outcomes contributing the most to the association detected. Here, we use the Variable Importance on Projection (VIP) defined for outcome *j* as VIP*_j_* = Cor(*Y_j_*, *Y*_PCEV_), where 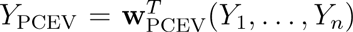. This VIP measure can be signed (as a correlation) or unsigned (in absolute value). In the case of PCEV-block, the VIP measure is defined in the same way, i.e correlation between the *original* outcomes and the final PCEV component.

### 2.5 Simulation study

Power and type I error of the PCEV approach are evaluated through extensive simulations assuming a sample size *n* = 500. In all simulations, a single continuous covariate *X* ~ *N*(0,1) is simulated and no confounder variables are included in the analysis. The multivariate outcomes **Y** are simulated under model (1) using a multivariate normal distribution of the residuals with variances equal to 1 and a block correlation structure made of five blocks of equal size. This correlation structure in **V***_R_* is governed by parameters *ρ_w_*, defined as the within-block correlation and *ρ_b_*, defined as the between-block correlation. The regression coefficients *B_k_* (*k* = 1,…, *p*), corresponding to the *k*th outcome, are chosen to match the values of *h*^2^ for each outcome, i.e. the proportion of variance explained by the covariate *X*, according to the relationship 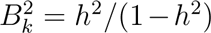. When multiple outcomes are associated with the variable *X*, we assume that each outcome explains the same proportion of variance *h*^2^.

We compare the PCEV approach with a PCR analysis and a PCEV-block approach. The PCR approach was selected for comparison because it does not involve any calibration (e.g. sparse methods typically require a tuning parameter) and because it provides a convenient framework for significance testing (which explains the exclusion of methods such as PLS or CCA, for example). In the PCEV approach, p-values are computed using the asymptotic test described above. In PCR, the first principal component of **Y** is extracted and then tested for association with the covariate. Finally, the PCEV-block approach is applied by using the same blocking structure as in the simulated data (i.e. five blocks). This analytical strategy is chosen to evaluate the performance of the PCEV-block approach in the situation where the mathematical property of independence between blocks is verified. Furthermore, we chose scenarios where the PCEV-block approach is not necessary (*p* ≪ *n*), allowing us to compare the performance of the PCEV-block with respect to the original PCEV.

We investigate the performance of the PCEV approach by varying several simulation parameters, namely *p* (the number of outcomes), *ρ_w_, ρ_b_* and *h*^2^ under five main scenarios described in Table 1. For each scenario, 500 simulated datasets were generated, and power was computed for *p* = 20, 50,100, 200, 300, 400, *ρ_w_* = 0, 0.5, 0.7 and *ρ_b_* = 0, 0.5, 0.7 with *ρ_w_* ≥ *ρ_b_*. Scenario 0 is designed to look at type I error rate and therefore none of the outcomes are associated with *X*. Scenarios 1 – 3 are designed to look at power under various settings. Finally, scenario 4 evaluates the robustness of the PCEV-block approach with respect to the independence of blocks assumption. In this case, PCEV-block is applied using three strategies to define the blocks: i) *b* = 5 blocks are used, where blocks coincide with the simulation design; ii) *b* = 10 blocks are used, where the two blocks in i) are each split in two; iii) *b* = 10 blocks are used, where blocks are chosen at random. These strategies are labeled PCEV_*b*1_, PCEV_*b2*_ and PCEV_*b*3_, respectively.

**Table 1:** Parameters used for the simulations. *p* represents the dimension of the outcome vector **Y**; *ρ_w_* and *ρ_b_* represent the within-block and between-block correlation in the simulated outcomes (with *ρ_b_* > *ρ_w_*); *h*^2^ represents the heritability of each outcome associated with *X*.

| Scenario | $h^2$ | Outcomes associated with $X$ |
| --- | --- | --- |
| 0 | 0 | None |
| 1 | 1% | 10 outcomes associated: $Y_1, \dots, Y_5$ and $Y_{p-4}, \dots, Y_p$ |
| 2 | 0.1% | 25% of the outcomes in first two blocks |
| 3 | 0.1% | 25% of the outcomes in all blocks |
| 4 | 0.1% | 25% of the outcomes in all blocks |

### 2.6 Datasets

All datasets used in this paper are publicly available, or available upon request. The bisulphite sequencing data is included in the R package pcev. The ARCTIC methylation data have been deposited in dbGAP under accession number [phs000779.v1.p1]. Finally, the ADNI data is archived in a secured and encrypted system provided by the LONI Image Data Archive (IDA). Applying for access to data requires the submission of an online application form.

#### Analysis of bisulfite sequencing data

The dataset contains measurements of DNA methylation levels derived from bisulphite sequencing around the BLK gene, located on chromosome 8. A total of 40 different samples are analysed from three cell types: B cells (8 samples), T cells (19 samples), and monocytes (13 samples). These samples are derived from whole blood collected on a cohort of healthy individuals from Sweden. Data were sequenced on the Illumina HiSeq2000 system. Missing values are imputed using the mean of neighbouring sites and the imputed data is available in the R package pcev. The region analysed contains 24,068 CpG sites, from which we removed the duplicate sites and analyzed the 25% most variable sites (which coincide with the sites having the largest depth), for a total of 5,986 sites spanning a region of 2.5 Mb. Methylation levels at each CpG sites are measured using the logit of the methylated proportions. To apply the PCEV-block approach, we took advantage of the natural clustering as a function of physical distance between CpG sites on the DNA strand. We clustered CpG sites such that sites within 500kb were grouped together. In order to obtain clusters with less than 30 sites (to ensure *p* ≪ *n*), we subsequently broke large clusters into smaller ones based on genomic distance. We obtained a total of 983 blocks of size ranging from 1 to 30 sites. Since this dataset is too large to use the classical PCEV method, we only used the PCEV-block framework to test the association between methylation levels and cell type (dichotomous variable, testing B-cells versus others).

#### Gene-based analysis of 450K Illumina methylation data

This analysis focuses on a sample of 1035 individuals who served as controls for the Assessment of Risk for Colorectal Tumors in Canada (ARCTIC) study [15, 16]. Methylation at 485,512 CpG sites was measured in stored lymphocyte samples using the Infinium Human Methylation450 BeadChip. We are interested in the association between methylation at the gene level and cigarette smoking status (which is dichotomous). By performing a gene-based analysis, we are lowering the epigenome-wide significance threshold, and therefore expect to see more associated genes. The CpG sites were allocated to 20,041 genes using the UCSC gene annotation; we also considered sites located 2kB upstream from transcription starting site and downstream of the 3’ end. With this definition, the genes under consideration contained anywhere between 2 and 288 CpGs, allowing us to perform PCEV using both the block and the classical approach since the number of CpGs per gene was substantially smaller than the number of individuals. Methylation data was normalised using Functional Normalisation [17], and further adjusted to account for cell type mixture using the reference-based method proposed by Houseman et al. [18] After normalisation, correction and exclusion of the X and Y chromosomes, we were left with 169,239 CpGs, located in 18,969 genes.

#### Analysis of brain imaging data

Data used in the preparation of this article were obtained from the Alzheimers Disease Neuroimaging Initiative (ADNI) database (adni.loni.usc.edu). The ADNI was launched in 2003 as a public-private partnership, led by Principal Investigator Michael W. Weiner, MD. The primary goal of ADNI has been to test whether serial magnetic resonance imaging (MRI), positron emission tomography (PET), other biological markers, and clinical and neuropsychological assessment can be combined to measure the progression of mild cognitive impairment (MCI) and early Alzheimers disease (AD).

The analysis we performed is based on data acquired from 340 participants from ADNI GO/2, for whom both genetic and PET data were available. [18F]Florbetapir PET imaging was employed to assess brain amyloid-*β* (A*β*) protein load using PET standardised uptake value ratios (SUVR) in 96 brain regions. These 96 A*β* levels represent the phenotypes of interest in our analysis. Association with diagnosis (Alzheimer versus others) was investigated. Phenotypes were adjusted for gender, age and education level directly within the PCEV framework.

## 3 Results

### 3.1 Simulation study

As one can see in Figure 1, Type I error for PCEV is well controlled and is not influenced by the number of variables nor by the correlation between outcomes. Here we have used the Wilks’ test for the classical PCEV, and performance is as expected even when the number of responses is as large as 400. Supplementary Table S1 shows that the Type I error for Wilks’ test is also well controlled at significance levels as low as 10^-4^.

**Figure 1:**
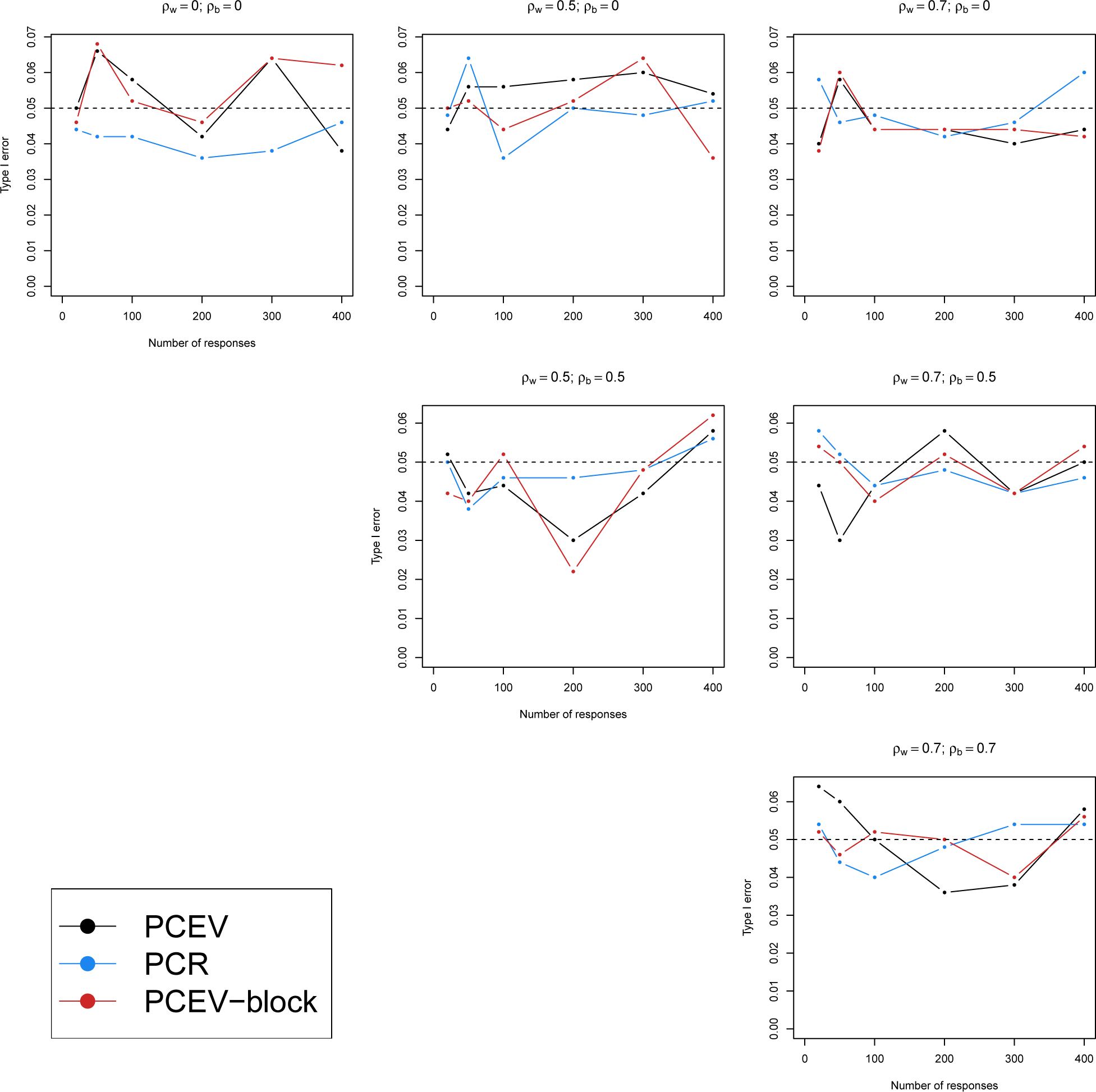
Type I error as a function of the correlation between outcomes *ρ* and the number of responses.

Figures 2, 3, and 4 illustrate the power of the PCEV, PCEV-block and PCR approaches in the next three simulated scenarios. In all cases, we observe that power of PCR is very low compared to the PCEV approaches. This is as expected, since PCR is not optimal with respect to identifying components maximally related to the covariate *X*. In all figures, it can also be seen that power increases with correlation between the outcomes. Such behaviour is also expected, since we can think of correlation as spreading out the signal across multiple outcomes, and it is therefore easier to detect it with the PCEV methods.

**Figure 2:**
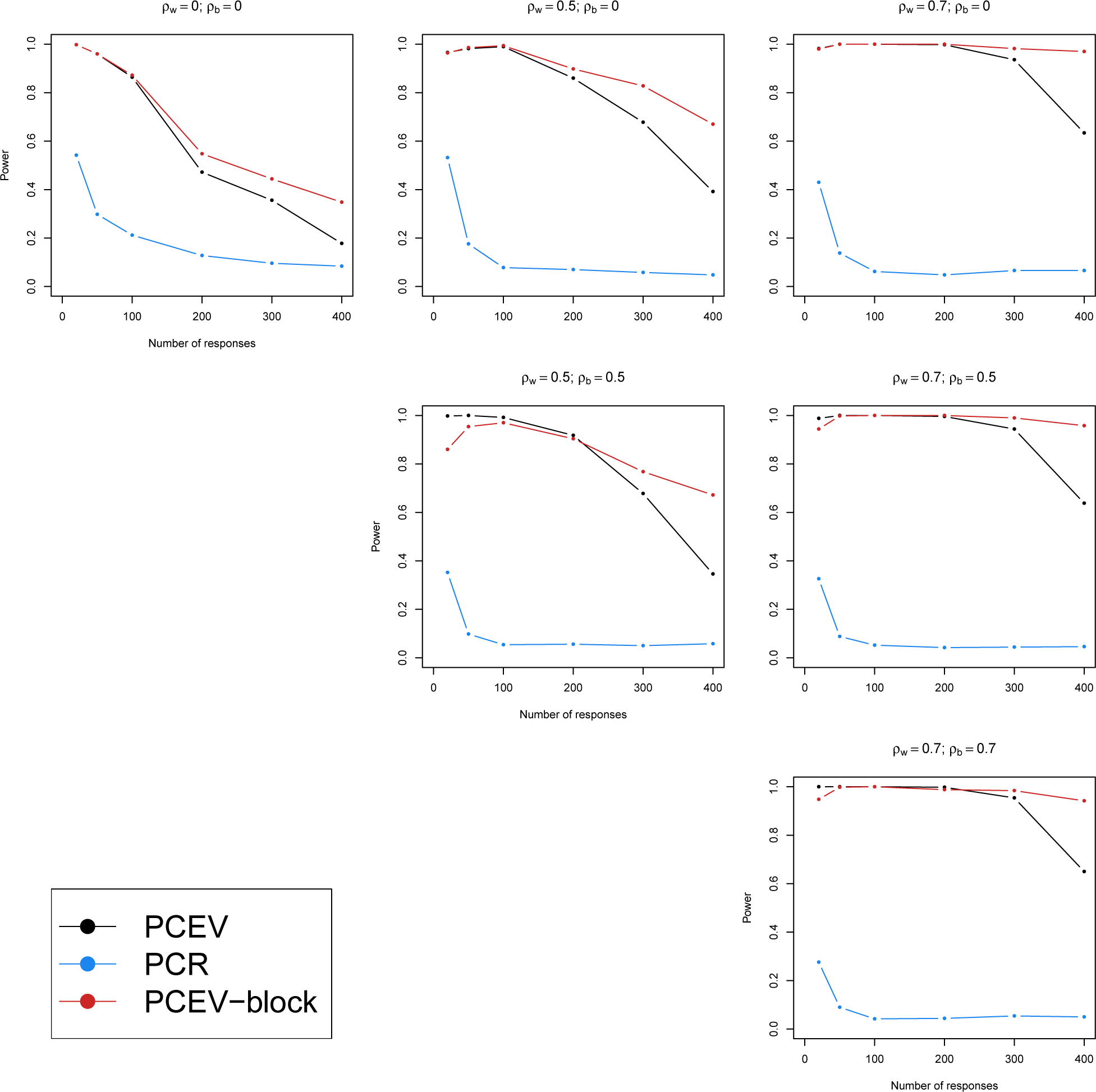
Power of PCEV: scenario 1.

**Figure 3:**
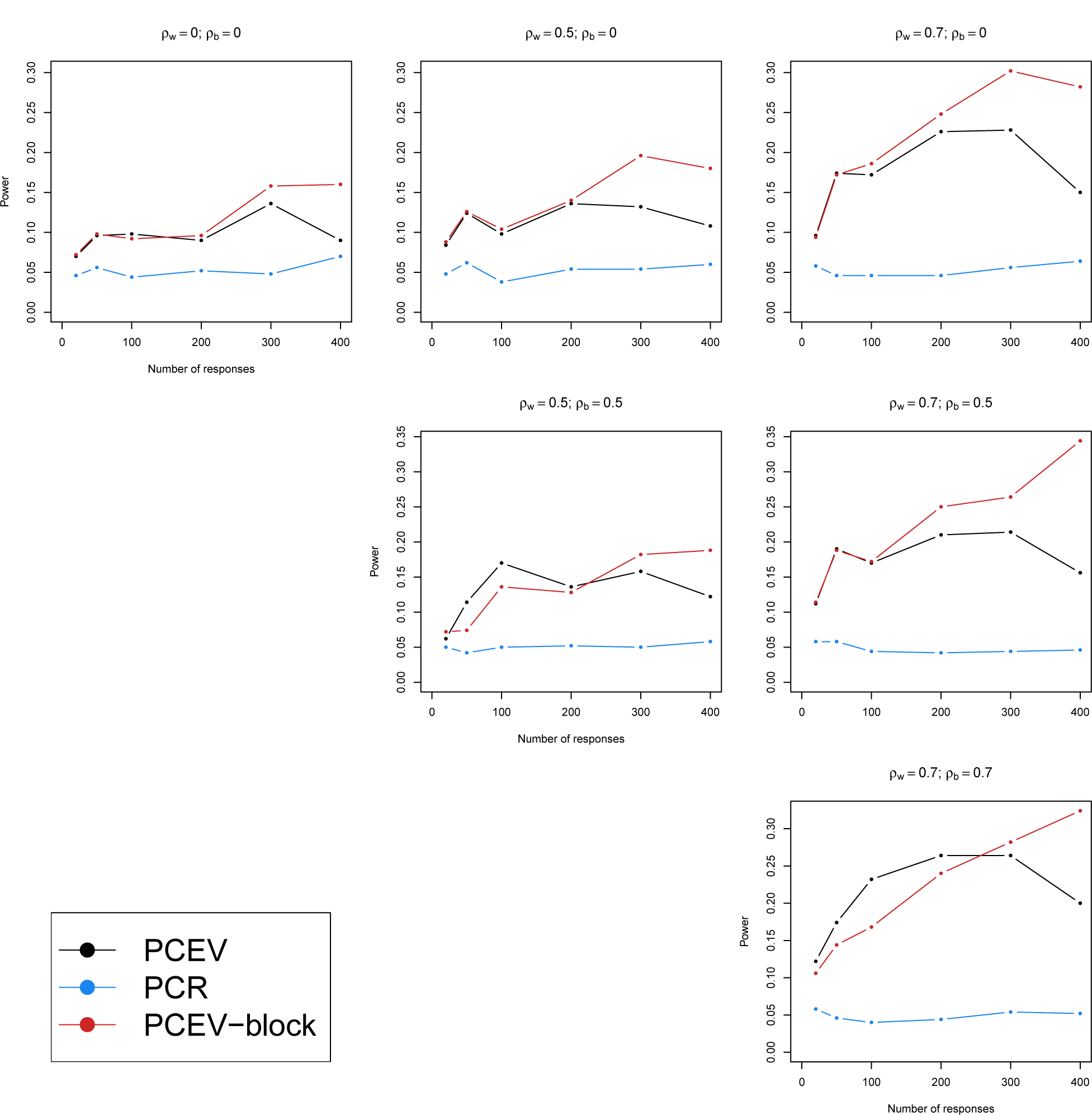
Power of PCEV: scenario 2.

**Figure 4:**
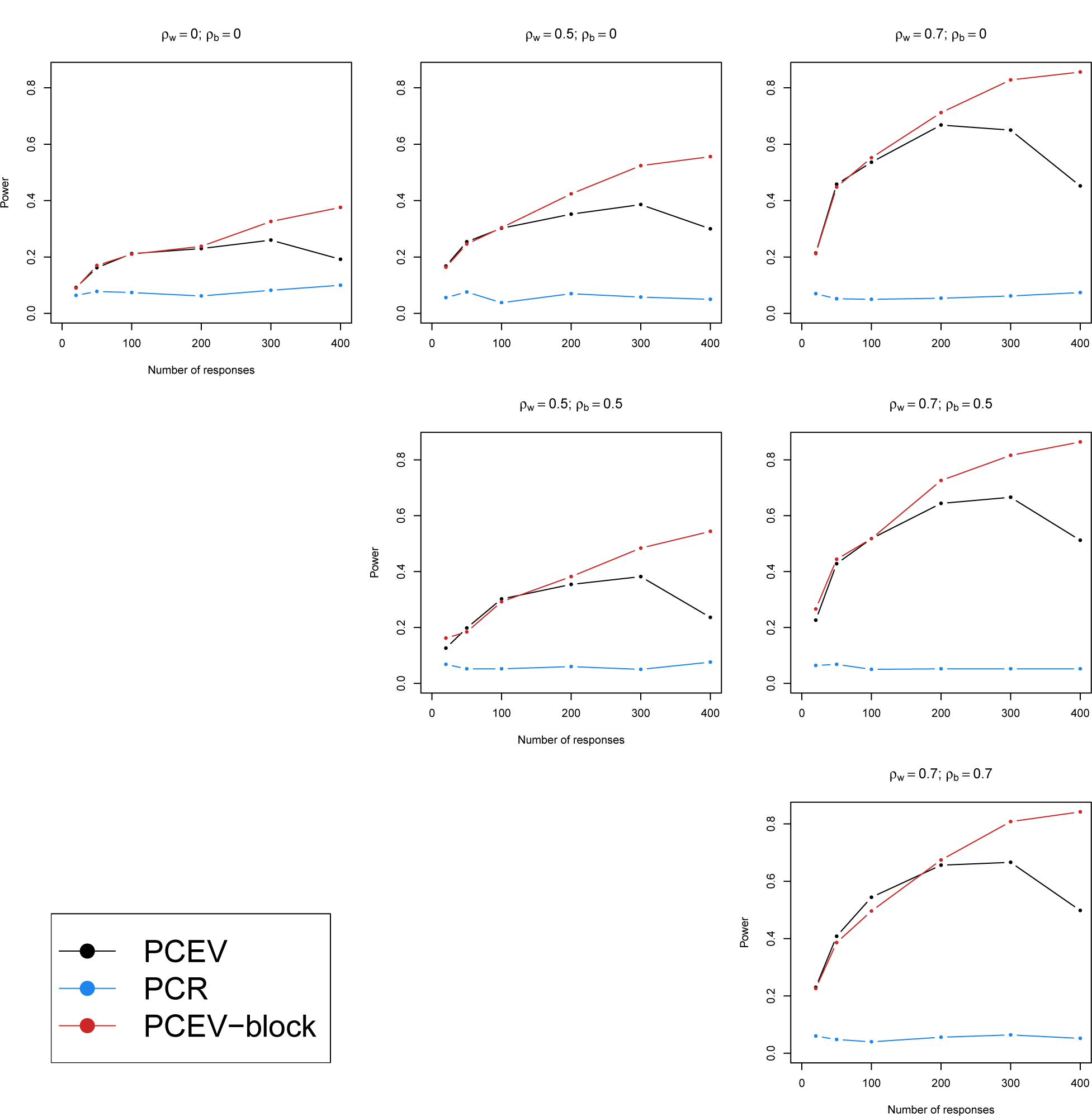
Power of PCEV: scenario 3.

However, the relationships between power and patterns of associated responses are more nuanced. Increasing the number of outcomes, i.e. increasing *p*, has a detrimental effect on power when the number of outcomes associated with *X* is low (scenario 1, Figure 2), and this is true for any of the methods that we explored. However, when the signal to noise ratio is kept constant as *p* increases (scenarios 2-3-4), power tends to increase with *p* up to a point, and then decrease afterwards. This inflection point seems to be where the number of outcome variables is large enough that the residual variance matrices become ill-conditioned. Once such a state is reached, the power of the PCEV approach decreases quickly (Figures 3 and 4) with larger values of *p*. However, crucially, the PCEV-block approach is not affected by this ill-conditioning phenomenon, since the residuals are estimated within each block. Therefore, each residual variance matrix is estimated from a much smaller number of outcomes. We also note that varying the between-block correlation *ρ_B_* in the simulation has a small impact on power, as it increases correlation between variables but does not spread the signal across outcomes. On the contrary, increasing the within-block correlation *ρ_W_* increases power, as expected.

Finally, Figure 5 illustrates the sensitivity of the PCEV-block approach with respect to how the blocks are chosen. As can be seen, power is reduced when the blocks are more correlated with each other. This is as expected, since the PCEV-block approach relies mathematically on the independence between blocks. However, as we will see in the data analysis section, despite the reduction in power, the VIP measures are very robust to misspecifications of the independence assumption.

**Figure 5:**
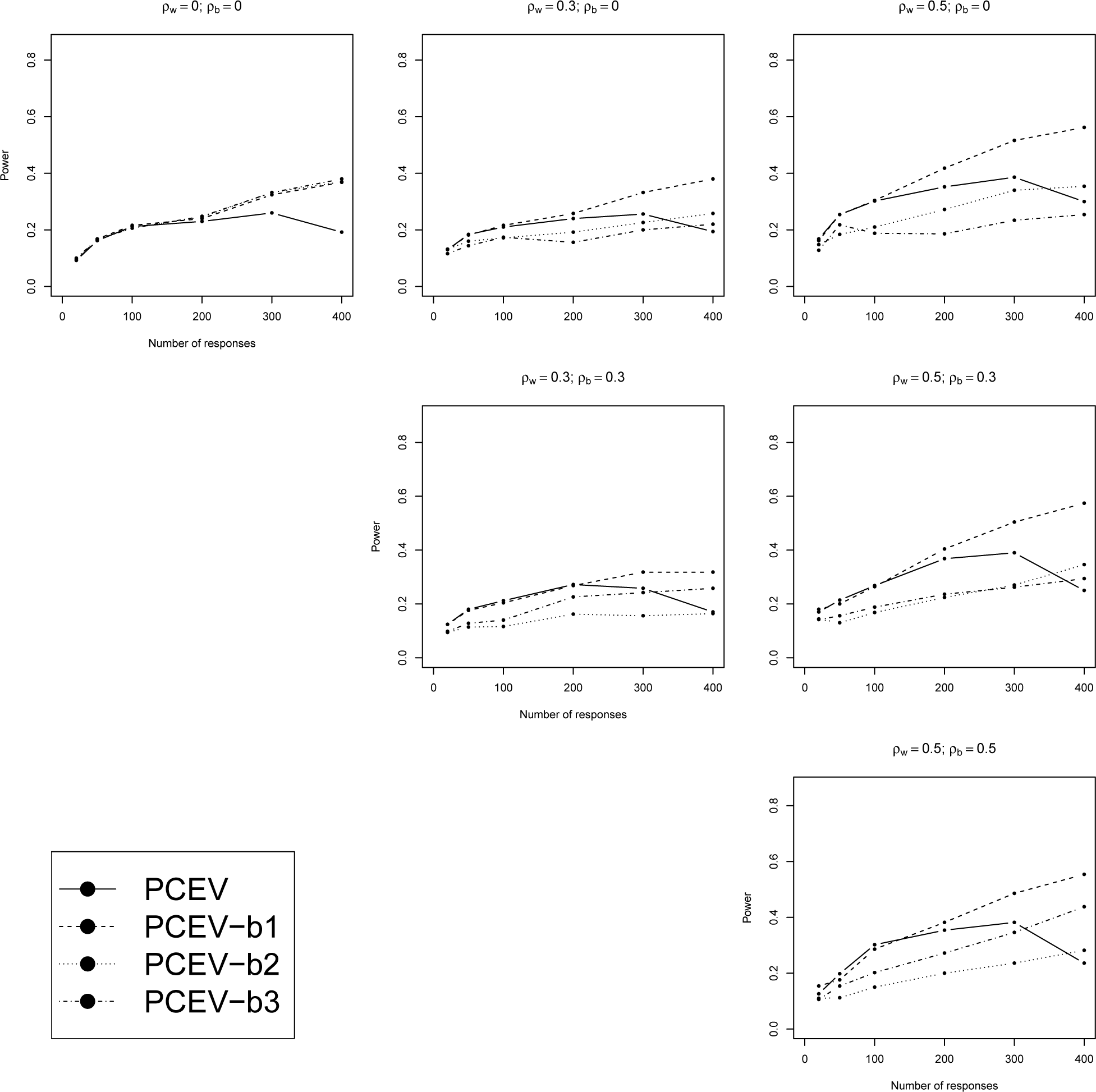
Power of PCEV: scenario 4

### 3.2 Data analysis

#### Analysis of bisulfite sequencing data

The genomic region we have analysed near the BLK gene is known to be hypomethylated in B-cells, compared to other cell types [19]. Since we have only 40 samples and over 5,986 sites in our region of interest, it is impossible to perform the regular PCEV test, and therefore we use the PCEV-block framework, and we expect to capture the regions association using only a single statistical test. Note that the methylation levels in this dataset show only a mild level of correlation, since 50% of the CpG sites pairs have a correlation smaller than 0.15 and 99% of the pairs have a correlation smaller than 0.50. As a result, the assumption of independence between blocks is not strongly violated in this example.

The p-value obtained with PCEV-block for all 5,986 sites simultaneously was 6 × 10^-5^, computed using 100,000 permutations. This result can be considered highly significant since only one test was performed. Furthermore, the results obtained are in excellent agreement with other approaches to analysis. For instance, Figure 6a shows the results using local linear regression to obtain a smoothed curve for each cell type in the region studied. As one can see, there is a region (delimited by vertical bars) comprising the BLK gene and a small upstream region where B-cells are differentially methylated compared to the two other cell types. Univariate regression analyses also confirmed this hypothesis (Supplementary Figure S1). Using the VIP measure, we are also able to identify which CpG sites contribute most to the association obtained (Figure 6b). Note that the region delimited in red in Figure 6b is identical to the region delimited by vertical bars in Figure 6a. Furthermore, there is a strong relationship between VIP measures and univariate p-values (Supplementary Figure S2, left panel). Signed VIP measures can also be used to detect the direction of the association, and we see in Supplementary Figure S2 (right panel) that such measures are highly correlated with the univariate slope regression coefficients.

**Figure 6:**
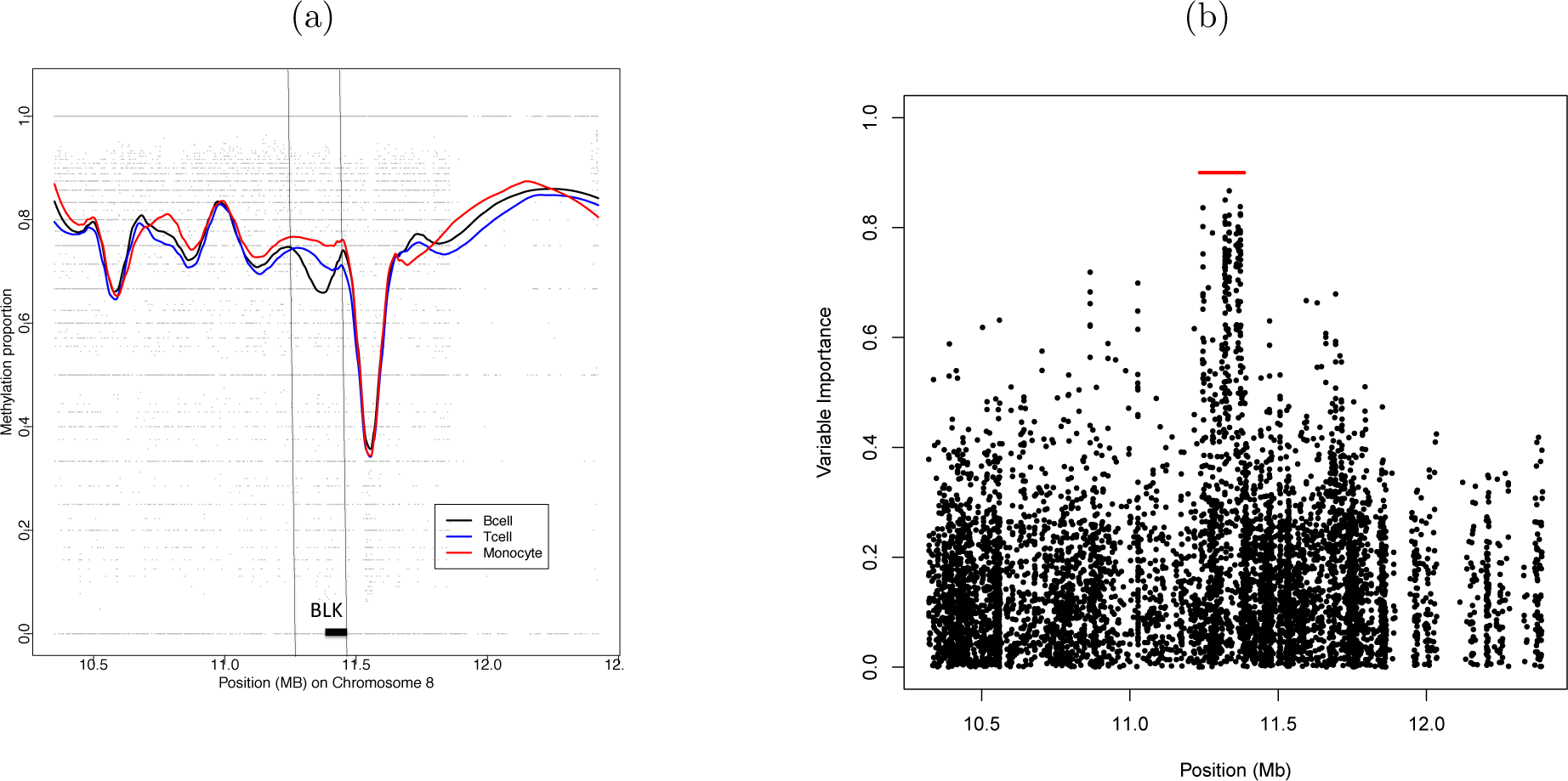
Methylation sequencing data: analysis of the BLK region, (a) Analysis using local linear regression: each curve represents the average loess smooth methylation predictions over cell types, (b) VIP (unsigned) measures for each CpG site.

#### Gene-based analysis of 450K Illumina methylation data

Analysis of the ARCTIC data was undertaken gene by gene, after selecting probes in or near each gene. In Figure 7, we show a scatter plot comparing the results of PCEV for each gene (without using the block version), and a gene level summary of the univariate p-values, where both are given on the negative log scale. The gene-level summary of the univariate p-values is defined as the minimum p-value among the CpGs tested in the gene, corrected for the number of independent CpG sites in the gene, i.e. the minimum p-value is multiplied by the estimated number of independent CpGs. The effective number of independent tests was estimated using the method proposed by Gao et al. [20] By examining the smallest p-values, or the top of this scatter plot, Figure 7 suggests that PCEV tends to enhance power in particular for the genes with the strongest association. In Figure 8, we compare the VIP values obtained from both the classical PCEV and the block approach, for four different genes F2RL3, AHRR, RARA, and GNG12 known to be associated with cigarette smoking [21, 22]. The PCEV-block approach proceeded by defining three blocks in each of these genes, and by allocating CpG sites to blocks in a linear fashion (for example, if a gene contained 30 sites, the first 10 were assigned to one block, the next 10 to a second block and so on). We purposely chose a non-optimal block definition so that we could evaluate the robustness of the block method to a poor choice of block (here, blocks were not chosen to be independent). As we can see, the VIP values provide information that is very similar to the univariate p-values across these four genes.

**Figure 7:**
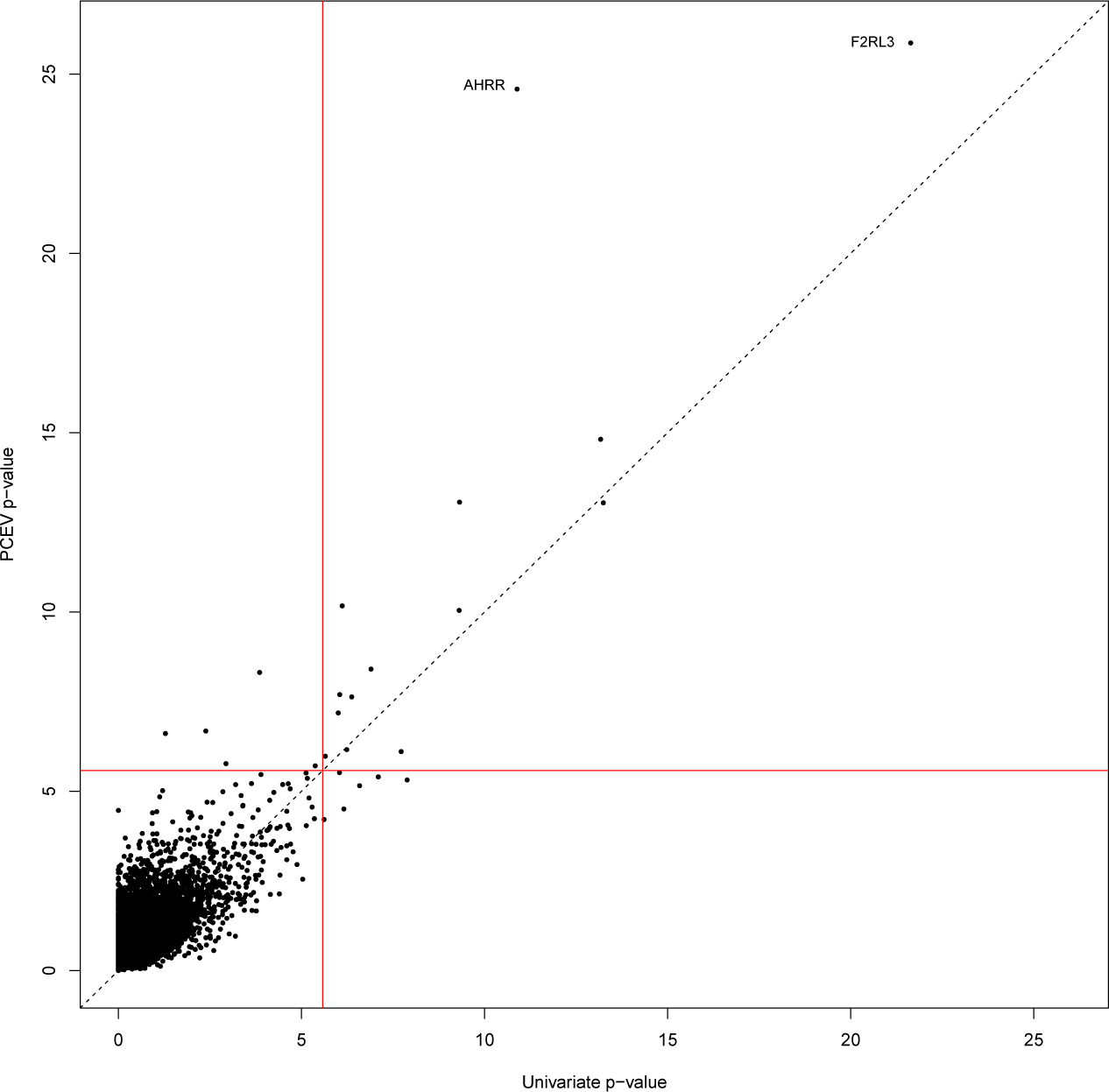
Comparison of the gene-based PCEV approach (without block) with a univariate analysis. Red lines correspond to significance threshold.

#### Analysis of brain imaging data

This dataset illustrates an example where *p* ≪ *n* and therefore no block strategy is necessary to perform a PCEV analysis. However, we performed both analyses (with and without the blocks) for the sake of comparison. In this data, there is a very high level of correlation between all brain regions, as shown in Supplementary Figure S3. Such high and extensive correlation makes clustering regions into blocks extremely challenging and we did not succeed in identifying well-separated cluster with any reasonable level of confidence. However, we present the results of the PCEV-block approach using 10 blocks obtained using a hierarchical clustering technique. Hence, we note that this dataset has several interesting features allowing us to i) compare results between PCEV and PCEV-block, and ii) evaluate the performance of the PCEV-block strategy, using the traditional PCEV approach as a gold standard, in a situation where clusters are very poorly chosen and highly correlated with each other. Analysis of the amyloid-beta accumulation against disease status (Alzheimer disease versus others) reveals a significant association regardless of the testing procedure chosen (see Table 2). Furthermore, Figure 9 illustrates the good agreement between variable importance measures computed using the classical PCEV approach and the block approach. We also note the monotonic relationship between univariate p-values (obtained from a simple regression analysis between amyloid-beta levels in each brain region and disease status) and VIPs.

**Figure 9:**
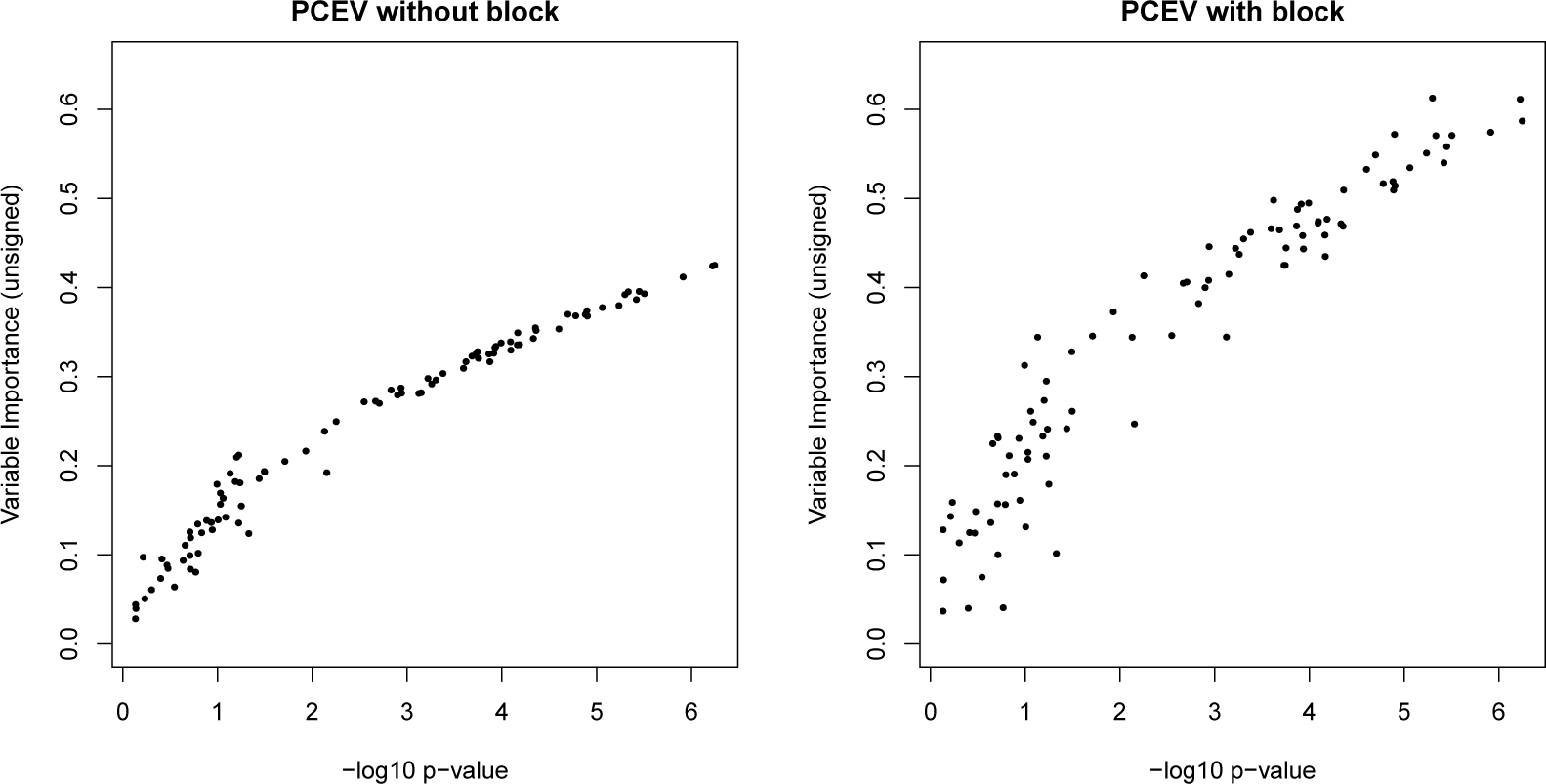
PCEV variable importance measures versus univariate p-values (negative log scale) for the association between amyloid-beta accumulation and disease status.

**Table 2:** P-values for the joint association between amyloid-beta accumulation and disease status. Permutation tests were performed using 100,000 permutations.

|  | PCEV | PCEV with blocks |
| --- | --- | --- |
| Exact test | $8.13 \times 10^{-5}$ | — |
| Permutation test | $2 \times 10^{-5}$ | $5 \times 10^{-5}$ |

## 4 Discussion

In this article, we have revisited a dimension-reduction approach, PCEV, that has unfortunately received little attention in statistical genetics or biostatistics in general. We showed how PCEV is well-suited for multivariate association studies, since it is optimal with respect to capturing the association between a multidimensional phenotype and a set of covariates.

We also presented a hypothesis-test framework which relies on an asymptotic result but no resampling. In particular, we presented two analytic tests that can be used with the traditional single-block PCEV: Wilks’ and Roy’s tests. The former should only be used when there is a single covariate and when *n* > *p* + 1, while the latter can be used in much more general settings (even when *p* ≫ *n*). Note also that Wilks’ test is exact, whereas Roy’s uses an asymptotic result and therefore is based on an approximation to the null distribution when the sample size is large enough. These asymptotic results mean that PCEV can be computationally extremely fast, which makes it feasible for use in large scale pipelines of multivariate analyses. For the block approach to PCEV, the independence of blocks assumption violates the distributional assumption necessary for Johnstone’s approximation [14] to be valid; for this reason, we have opted for a permutation procedure in order to test for association. Finally, although our discussion has focused on testing the first PCEV component (which is the only one computable when *X* is a single covariate), this framework can also be applied to several PCEV components independently, when multiple covariates are analysed simultaneously.

We compared the performance of PCEV to PCR, which is a very popular approach for region-based analyses [23, 24, 25, 26, 27, 28, 29]. Our simulation results show that, as can be expected, there is no guarantee that the first principal component is at all associated with the covariates. PCR had very low power to detect association, especially when compared to both the classical and block approaches to PCEV. Hence, in general, we discourage the use of PCR for region-based analyses of multidimensional phenotypes. We note that our focus was restricted to computationally efficient dimension reduction approaches with an asymptotic or exact testing framework, and therefore, we have purposely ignored methods that require tuning parameters or rely entirely on bootstrap or permutation strategies to perform inference. For this reason, we did not compare our proposed block approach to other regularization approaches to PCEV [5, 7], and furthermore we did not compare to regularized and sparse forms of CCA and PLS. However, we show that there are strong statistical connections between PCEV and other component-based methods such as MANOVA, PLS, CCA, and LDA (details are given in Appendix 4). For example, we show that PCEV also looks implicitely for an optimal linear combination of the covariates **X**, with maximal correlation with YPCEV.

Our analysis of the bisulfite sequencing data reveals how useful the block approach can be in the presence of high-dimensional data. Indeed, most multivariate methods do not give meaningful results for such a dataset, due to the discrepancy between the sample size and the number of variables. In contrast, the block PCEV approach was able to recapitulate known results about the potential existence of a differentially methylated region (DMR) around the BLK gene. As shown in Figure 6b (right panel), PCEV-block captures the association between 5,986 variables and the cell-type covariate in a single test, despite the fact that there are only 40 subjects. Moreover, the variance importance factors (VIP) were able to accurately pinpoint the source of the signal among the 6,000 CpG sites (Figure 6a). Supplementary Figure S2 (left panel) shows that VIP correlates well with nominal p-values from univariate tests and Supplementary Figure S2 (right panel) indicates that signed-VIP values correlate with univariate regression coefficients measuring the association between each variable and cell type.

With the ARCTIC dataset, our aim was to confirm the association results shown in Breitlin et al. [21] between methylation and cigarette smoking. Two genes showed particularly strong associations in Figure 7, F2RL3 (PCEV p-value: ≈ 10^-26^), as well as AHRR (PCEV p-value: ≈ 10^-25^), and both are previously reported smoking meQTL loci [21, 22]. By pooling all CpG sites at or near each single gene, our power to detect this associations was greater than what we could have achieved with an adjusted univariate approach. After using the VIPs to decompose the multivariate signals at these two genes (Figure 8), it is evident that for F2RL3, the signal is mostly driven by one CpG site, whereas for AHRR, there seem to be five sites contributing to the overall signal. In the latter case, this “pooling of forces” probably leads to the power increase.

**Figure 8:**
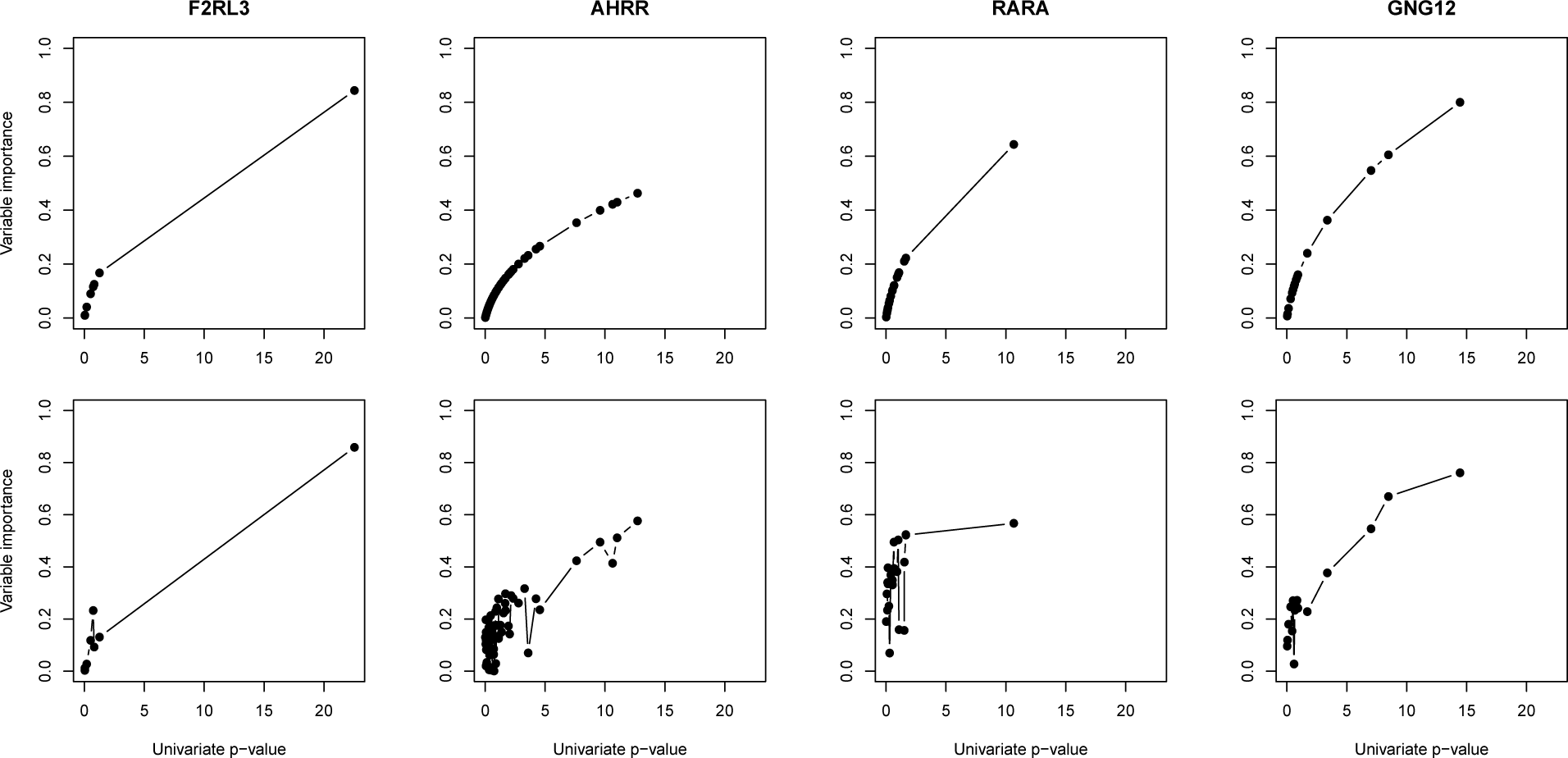
Comparison of the variable importance (VIP) measures for four genes, F2RL3, AHRR, RARA, and GNG12, known to be associated with cigarette smoking. For each gene, the top panel shows the VIP obtained from the PCEV without block, while the bottom panel shows the VIP obtained from the PCEV with block.

In the context of the brain imaging study, we were able to investigate the effect of wideranging correlations on the block approach to PCEV. We showed that, even though the independence of block assumption was clearly violated, the p-value obtained using permutations was comparable to that obtained from a classical approach to PCEV (i.e. without blocks) and an exact test. Furthermore, the VIP factors were similar for both approaches. Therefore, we see that the block approach is quite robust to violations of the assumption.

In summary, we have showed how PCEV is particularly well-suited for finding multivariate signals with widespread association with a set of covariates. There is a fine balance to strike when performing a multivariate association test with a very high-dimensional phenotype: although we want to include multiple correlated response variables and thus borrow strength across phenotypes, at the same time we would like to retain interpretability of the multivariate signal. Suppose, to give an extreme example, that in the context of a genome-wide methylation association study, an analyst decided to include all the CpG sites available on a microarray as an enormous set of outcomes, and then to test for global association, at the genome level, with a covariate. Rejecting the null hypothesis in such a scenario would be of no help whatsoever in targeting the source of this signal. This example also illustrates the fact that PCEV is not a variable selection method, and our simulations showed that when the signal is very sparse and the signal-to-noise ratio is low, power is greatly reduced. In contrast, if the analyst decides to severely limit the number of outcomes jointly analyzed, then targeting the source of any identified associations will be straightforward, but there will be little benefit over univariate testing. In general, choosing an appropriate set of outcomes for joint analysis, i.e. finding the right balance between interpretability and power, is an important consideration that should consider the context and the biology. We feel that region-based analyses, where regions could be parts of the brain, genes, pathways or some other external data partitioning, is where this method should be considered and can shine.

## 5 Software

An R package called pcev, implementing both the block and the classical approach to PCEV, is currently available:

- on CRAN (https://cran.r-project.org/web/packages/pcev/);
- and on GitHub (https://github.com/GreenwoodLab/pcev).

## Acknowledgements

Data collection and sharing for this project was funded by the Alzheimer’s Disease Neuroimaging Initiative (ADNI) (National Institutes of Health Grant U01 AG024904) and DOD ADNI (Department of Defense award number W81XWH-12-2-0012). ADNI is funded by the National Institute on Aging, the National Institute of Biomedical Imaging and Bioengineering, and through generous contributions from the following: AbbVie, Alzheimers Association; Alzheimers Drug Discovery Foundation; Araclon Biotech; BioClinica, Inc.; Biogen; Bristol-Myers Squibb Company; CereSpir, Inc.; Eisai Inc.; Elan Pharmaceuticals, Inc.; Eli Lilly and Company; EuroImmun; F. Hoffmann-La Roche Ltd and its affiliated company Genentech, Inc.; Fujirebio; GE Healthcare; IXICO Ltd.; Janssen Alzheimer Immunotherapy Research & Development, LLC.; Johnson & Johnson Pharmaceutical Research & Development LLC.; Lumosity; Lundbeck; Merck & Co., Inc.; Meso Scale Diagnostics, LLC.; NeuroRx Research; Neurotrack Technologies; Novartis Pharmaceuticals Corporation; Pfizer Inc.; Piramal Imaging; Servier; Takeda Pharmaceutical Company; and Transition Therapeutics. The Canadian Institutes of Health Research is providing funds to support ADNI clinical sites in Canada. Private sector contributions are facilitated by the Foundation for the National Institutes of Health (www.fnih.org). The grantee organization is the Northern California Institute for Research and Education, and the study is coordinated by the Alzheimer’s Disease Cooperative Study at the University of California, San Diego. ADNI data are disseminated by the Laboratory for Neuro Imaging at the University of Southern California.

This work was also supported by the Natural Sciences and Engineering Research Council of Canada (AL), the Fonds de recherche Sante (chercheur boursier, AL) and Nature et Technologies (Doctoral award, MT) du Quebec. The authors acknowledge the Ludmer Centre for Neuroinformatics and Mental Health for financial support, and they also thank Dr. Tomi Pastinen for providing the bisulfite sequencing data.

## Appendix

### Appendix 1: proof of Theorem 1

The proof of Theorem 1 will follow from the proof of the following, more general result.

**Theorem 2**. *Let* **A** *and* **B** *be two positive semidefinite matrices of dimension p × p, with* **B** *invertible. Assume* **B** *is block diagonal:*

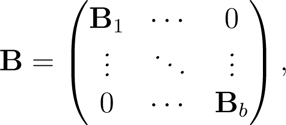

*and decompose A similarly:*

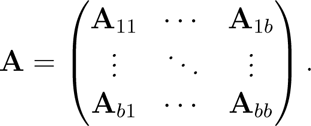

*Then the maximisation of the expression*

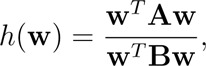

*can be decomposed in b* + 1 *distinct maximisations, as follows:*

*1. Maximise the expression*

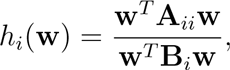

*for i* = 1,…, *b; let* **u***_i_ be the solution of the i-th maximisation.*

*2. Define the following matrices:*

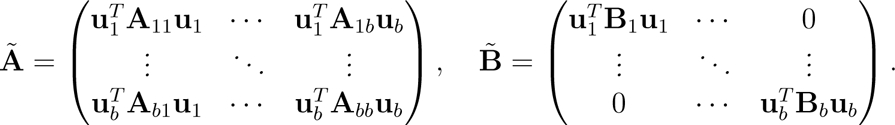

*Maximise the expression*

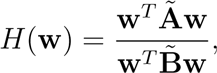

*and denote the maximiser of H* (**w**) *by* **v** = (*υ*1,…, *υ_b_*).

*3. The maximiser of the original expression h*(**w**) *is thus given by* **w** = (*υ*_1_ **u**_1_,…, *υ_b_* **u***_b_*).

*Proof*. We start by solving the original maximisation problem. Let **L** be a square-root of **B**, so that **B** = **LL***^T^*. We thus have

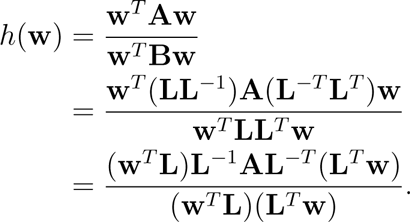

Therefore, if c is an eigenvector of **L**^-1^**AL**^-*T*^ associated to its largest eigenvalue, then **w**:= **L**^-*T*^**c** maximises *h*(**w**).

Since **B** (and thus **L**) is block diagonal, we can be explicit about the matrix **L**^-1^ **AL**_-*T*_; namely, we have

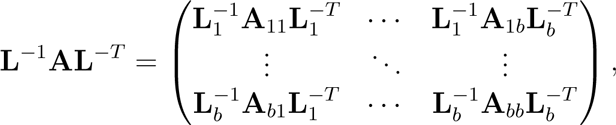

where **L***_j_* is the *i*-th block of **L**.

Now, we repeat the same analysis but with the first *b* maximisations. For a fixed *i*, we have

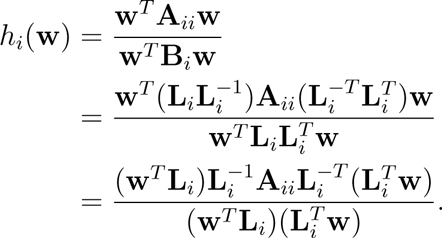

Similarly as above, if **c***_i_* is an eigenvector of 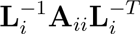 associated to its largest eigenvalue, then 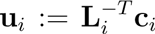 maximises *h_i_* (**w**); let **u** = (**u**_1_,…, **u***_b_*) be the concatenation of all these vectors, and note that its length is equal to *p*.

The final maximisation takes more care. First, we note that

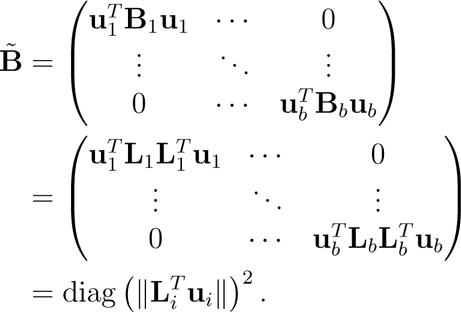

We have thus found a square root for the *b* × *b* diagonal matrix 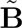.

As above, the maximisation of *H*(w) is related to an eigenvalue problem. In this case, we are looking for an eigenvector of the matrix

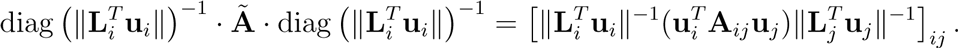

Let **d** = (*d*_1_,…, *d_b_*) be an eigenvector of this matrix associated to its largest eigenvalue. It finally follows that

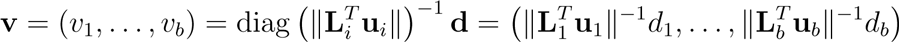

maximises *H*(w).

Recall our claim that **w** = (*υ*_1_ **u**_1_,…, *υ_b_* **u**_*b*_) maximises the original criterion

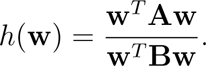

We can write

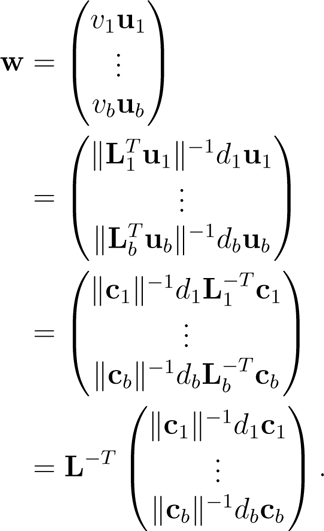

Therefore, to finish the proof, we need to show that 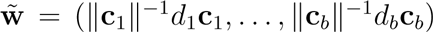 is an eigenvector of **L^-1^AL_-*T*_** associated to its largest eigenvalue. In other words, we need to show that 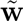 maximises the ratio

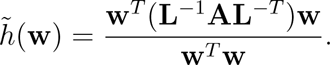

We first compute the denominator:

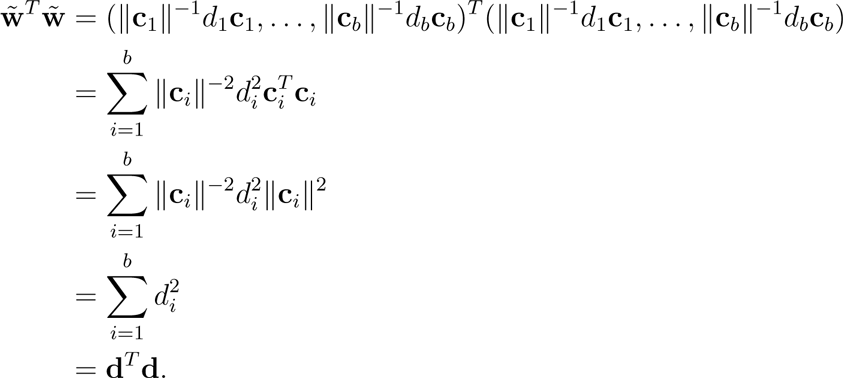

On the other hand, the numerator is given by

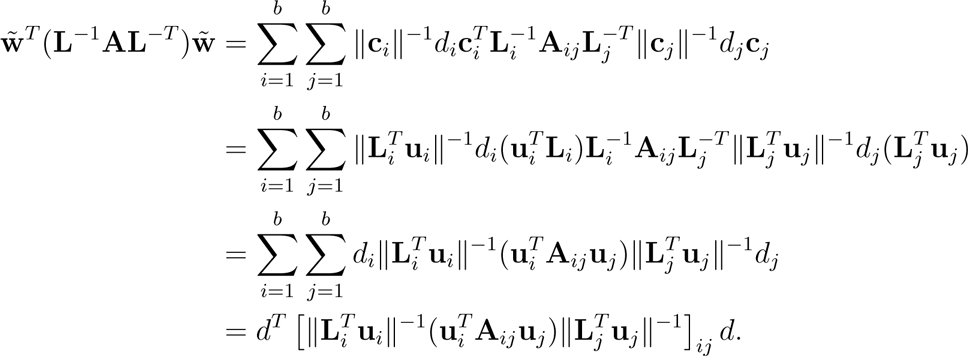

In other words, the ratio 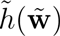 evaluated at 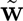 is equal to

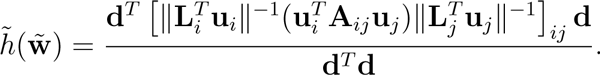

But since **d** is an eigenvector for the matrix 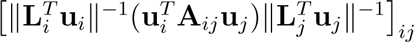 associated to its largest eigenvalue, we know that this ratio is maximised. This concludes the proof.

As a consequence of this theorem, we have proven the validity of the block approach. Note that since the maximisation of the two ratios

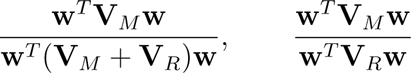

is equivalent, it follows that the independence assumption needed for the block approach can be made either conditional on the covariates **X** or unconditionally. Moreover, because of the generality of the theorem, we can deduce that the block approach also works for extracting further components; one simply needs to regress the first component on **Y** and perform PCEV on the residuals to get the second component, and so on.

### Appendix 2: Wilks test of significance

Let *X* be a single covariate, and consider the multivariate regression of *X* on a *p*-dimensional response vector **Y** = (*Y*_1_,…, *Y_p_*). It is well known that

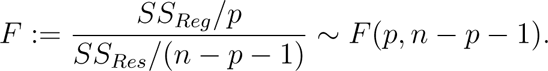

Moreover, note that we can rewrite *F* as follows:

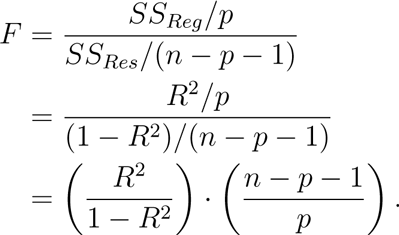

It remains to show that λ = *R*^2^/(1 – *R*^2^). But since *R*^2^ is the coefficient of multivariate correlation between *X* and the *p* variables *Y*_1_,…, *Y_p_*, *R*^2^ is defined as the maximum correlation between *X* and any linear combination of **Y:**

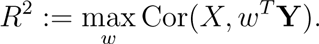

Therefore, *R*^2^ must be equal to the first canonical correlation between **Y** and *X*. Hence, we have

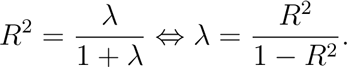

This completes the proof.

### Appendix 3: Roy’s largest root test statistic

In the hypothesis test framework developed above, we presented two tests based on the largest eigenvalue λ of the matrix 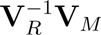, which is also the largest root of the determinantal equation

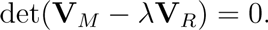

Under the assumption that there is only one covariate, we can derive the exact distribution of the ratio λ/(1 + λ); as noted in Appendix 2, this results holds as long as *n* > *p* + 1, where n is the sample size and *p*, the number of response variables.

More generally, Johnstone [30] derived an approximation to the distribution of λ, after a suitable transformation. The result, using the notation of this article, is given below:

**Theorem 3**. *[14] As n, q, p → ∞, we have*

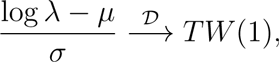

*where TW* (1) *is the Tracy-Widom distribution of order 1 [31], and μ, σ are defined as follows: let s* = min(*p, q*) *and write*

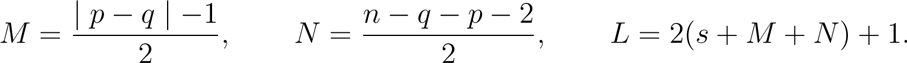

*Next, define*

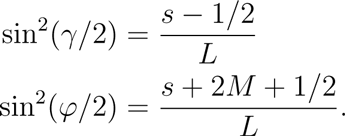

*Finally, we set*

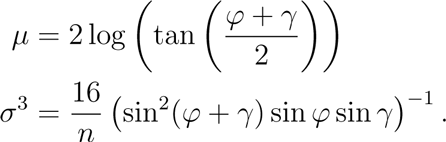

Under the scenario where there is only one covariate and *n* > *p* + 1, we recommend using the Wilks’ Lambda test, since it typically has higher power than the test based on Johnstone’s approximation. In any case, Theorem 3 can be used in much more generality, even when *p* ≫ *n*.

### Appendix 4: Relationship to other multivariate approaches

#### Notations

To discuss the relationship of PCEV with other existing methods, one needs to introduce some notations and background. Assume that the vectors *Y* and *X* are measured on the same sampling unit *i*, (*i* = 1,…, *n*), and denote **Y** and **X** the matrices *n* × *p* and *n* × *q* of the observed data, respectively. Without loss of generality, we will assume that **Y** and **X** are centred matrices. Model (1) in the article leads to a multivariate regression model between **Y** and **X**. Thus, one can write

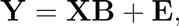

where the matrix **B** is the coefficients of the model.

The least square estimator of **B** is given by

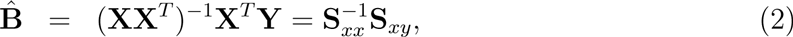

where S*_xx_* and S*_xy_* are block forms of the overall sample covariance matrix **S** of the vector (*y*_1_,…, *y_p_*, *x*_1_,…, *x_q_*), with **S** given as

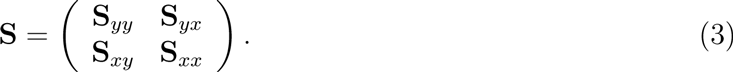

One can also write the total sum of squares and products as

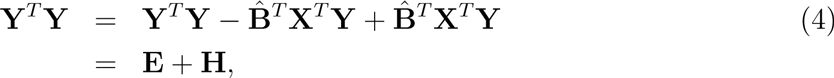

where

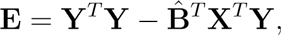

is the overall sum of squares and products that is unexplained by **X** (i. e. residuals) and

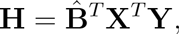

is the overall regression sum of squares and products matrix (i. e. variance explained by **X**). Notice also that following (2), one can write **E** and **H** in terms of the blocks form of the overall sample covariance matrix **S** in (3) as follow:

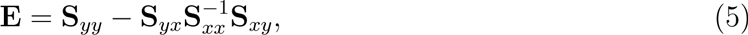

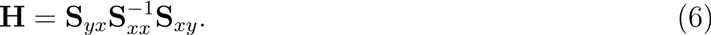

#### PCEV and its relationship to CCA

Recall that PCEV seeks a linear combination of outcomes, **w^T^Y**, which maximises the ratio *h*^2^(**w**) of variance being explained by **X**. Thus, following (4), the heritability *h*^2^(**w**) defined in section 2.1 of the paper can be written as

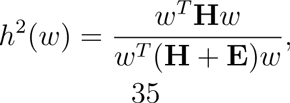

If one substitutes **E** and **H** by their values from equations (5) and (6), then one can write

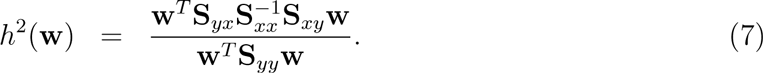

Thus, the vector **w** that maximises *h*^2^(**w**) in (7) can be obtained as the first eigenvector of 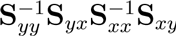, which is the first canonical direction on **Y**. The maximum heritability in this case is the largest squared canonical correlation, 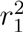.

Therefore, even if PCEV seeks the best linear combination of **Y**, since it maximises the ratio of the variance explained to total variance, it implicitly looks for the best linear combination of **X** with maximum correlation with 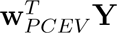.

#### PCEV and its relationship to LDA and one-way MANOVA

Assume that we have *K* groups (e. g. factor with *K* levels) and assume that the *p* × 1 vector **Y** is measured on the sampling unit *i*, (*i* = 1,…, *n*) for each group *k* = 1,…, *K*, and denote **Y** the matrices *nK* × *p* of the observed data. Notice that here we deal with a simple balanced case where *n* the number of units at each group is the same. All the results remain the same for the unbalanced case.

In order to establish a relationship between PCEV and LDA (and one-way MANOVA), one needs to rewrite the MANOVA model as a multivariate regression model. To do so, one needs to assume first *K* – 1 dummy variables *x_k_, k* = 1…, (*K* – 1), defined as

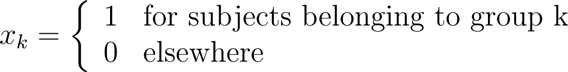

Then, one can write the MANOVA model as the multivariate regression of **Y** on the dummy variables *x_k_, k* = 1…, (*K* – 1). One can write,

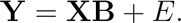

Following equation (4), one can verify that **E** and **H** are exactly the within sum of squares and products and the between sum of squares and products, respectively.

Again, from the definition of heritability, one can see that PCEV searches for direction on the column space of **Y** that maximises the ratio of the variation explained by the groups to the total variation. That is, one can write

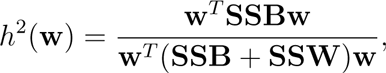

where **SSB** and **SSW** are the within sum of squares and products and the between sum of squares and products, respectively. Thus, the vector *w_PCEV_* that maximises the heritability is exactly the first linear discriminant function. In the MANOVA context, *h*^2^(**W***_PCEV_*) is termed *Roy’s largest root test*. Another interpretation of *h*^2^(**W***_PCEV_*) is as the maximum squared correlation 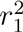, between the first discriminant function and the best linear combination of the *K* – 1 dummy variables. The later interpretation can be concluded from the relationship between PCEV and CCA.

Note that the heritability or *Roy’s largest root test* statistic is equal to

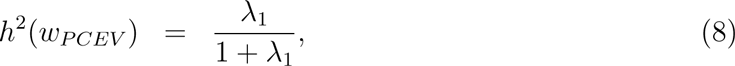

where λ_1_ is the largest eigenvalue of **SSW^-1^SSB**. For *K* > 2, its distribution is difficult to determine but an approximation can be used to calculate an exact p-value. SAS uses Davies approximation to calculate the p-value. In the case of only two groups (*K* = 2), λ_1_ is proportional to the Hoteling statistic which is the generalisation of the squared of the *t*-test statistic in the univariate case. Thus λ_1_ can be transformed to a Fischer exact-test statistic, and can be used instead of *h*^2^(**W**_PCEV_).

Finally, we note that the connection between LDA and PLS has been described before [32].

### Appendix 5: Supplementary figures and results

**Table S1:** Table S1: Type I error as a function of the within-block correlation *ρ_W_* and the number of responses *p*, for several significance levels.

| Within-block correlation | Significance level | Number of response variables |  |  |  |  |  |
| --- | --- | --- | --- | --- | --- | --- | --- |
| | | $p = 20$ | $p = 50$ | $p = 100$ | $p = 200$ | $p = 300$ | $p = 400$ |
| $\rho_W = 0$ | 0.1 | 0.09922 | 0.09978 | 0.09958 | 0.10139 | 0.09971 | 0.09954 |
|  | 0.05 | 0.05044 | 0.04908 | 0.04996 | 0.05057 | 0.05033 | 0.05089 |
|  | 0.01 | 0.01015 | 0.00994 | 0.01004 | 0.01008 | 0.00970 | 0.01030 |
|  | 0.005 | 0.00550 | 0.00498 | 0.00512 | 0.00503 | 0.00473 | 0.00530 |
|  | 0.001 | 0.00108 | 0.00108 | 0.00110 | 0.00084 | 0.00096 | 0.00109 |
|  | 0.0005 | 0.00055 | 0.00058 | 0.00050 | 0.00033 | 0.00053 | 0.00069 |
|  | 0.0001 | 0.00007 | 0.00011 | 0.00011 | 0.00005 | 0.00009 | 0.00016 |
| $\rho_W = 0.5$ | 0.1 | 0.09902 | 0.10070 | 0.09958 | 0.10139 | 0.09971 | 0.09954 |
|  | 0.05 | 0.04960 | 0.05010 | 0.04996 | 0.05057 | 0.05033 | 0.05089 |
|  | 0.01 | 0.00993 | 0.00979 | 0.01004 | 0.01008 | 0.00970 | 0.01030 |
|  | 0.005 | 0.00497 | 0.00495 | 0.00512 | 0.00503 | 0.00473 | 0.00530 |
|  | 0.001 | 0.00093 | 0.00078 | 0.00110 | 0.00084 | 0.00096 | 0.00109 |
|  | 0.0005 | 0.00043 | 0.00043 | 0.00050 | 0.00033 | 0.00053 | 0.00069 |
|  | 0.0001 | 0.00009 | 0.00011 | 0.00011 | 0.00005 | 0.00009 | 0.00016 |
| $\rho_W = 0.7$ | 0.1 | 0.09977 | 0.10118 | 0.09943 | 0.09863 | 0.10001 | 0.09977 |
|  | 0.05 | 0.05022 | 0.05105 | 0.04874 | 0.04915 | 0.04981 | 0.05036 |
|  | 0.01 | 0.01005 | 0.00994 | 0.00974 | 0.01022 | 0.00952 | 0.01013 |
|  | 0.005 | 0.00490 | 0.00504 | 0.00485 | 0.00532 | 0.00462 | 0.00490 |
|  | 0.001 | 0.00109 | 0.00108 | 0.00101 | 0.00113 | 0.00094 | 0.00080 |
|  | 0.0005 | 0.00049 | 0.00064 | 0.00054 | 0.00064 | 0.00039 | 0.00036 |
|  | 0.0001 | 0.00013 | 0.00011 | 0.00012 | 0.00010 | 0.00013 | 0.00010 |

**Figure S1:**
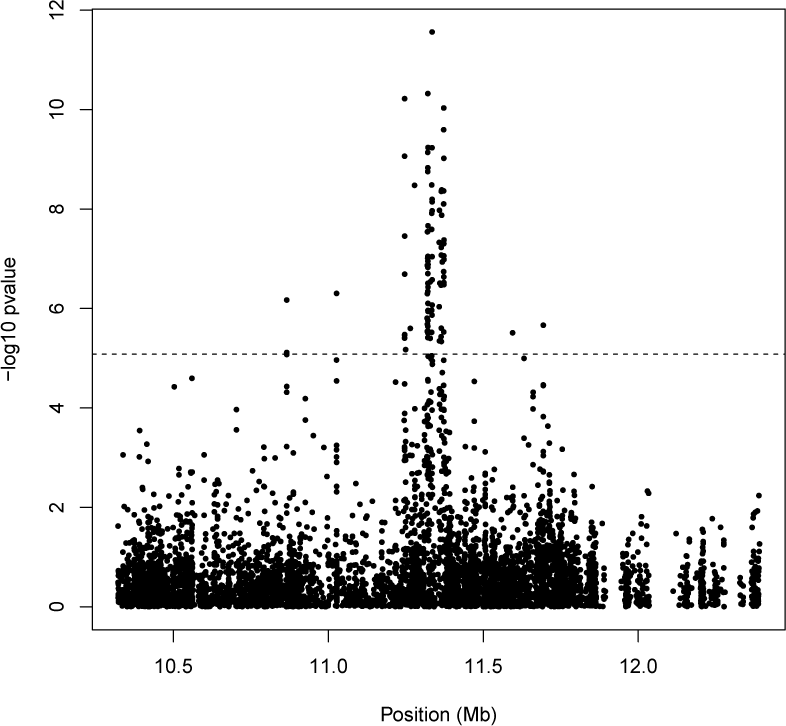
Methylation sequencing data: univariate analysis of the BLK region

**Figure S2:**
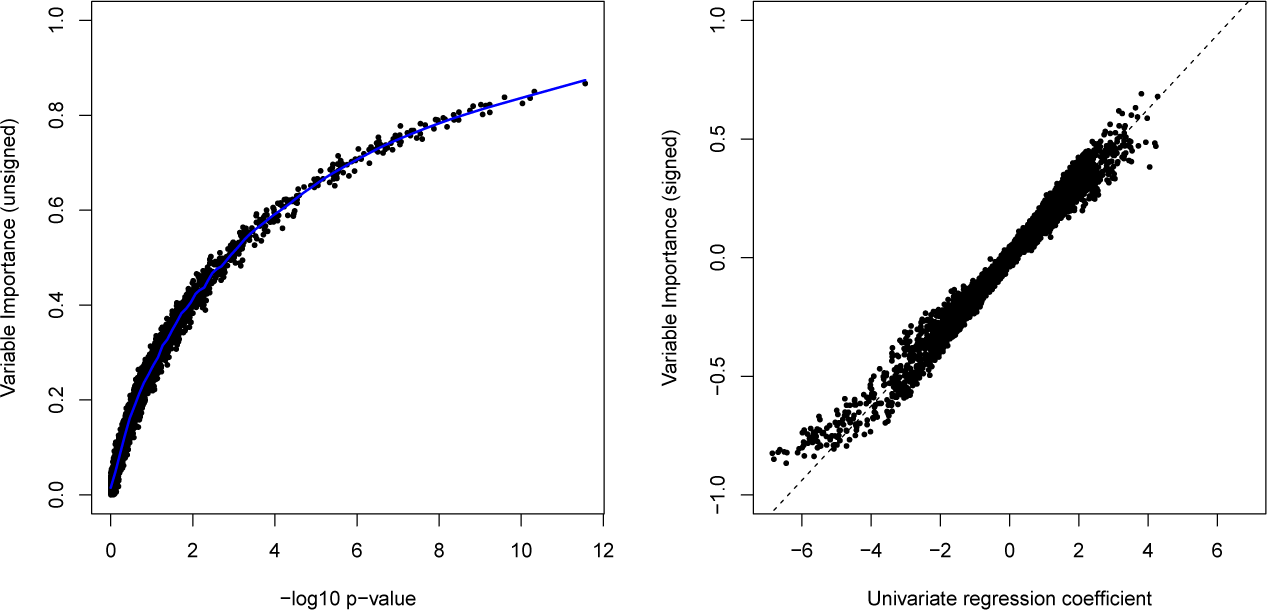
Methylation sequencing data: relationship between VIP, univariate regression coefficients and univariate p-values

**Figure S3:**
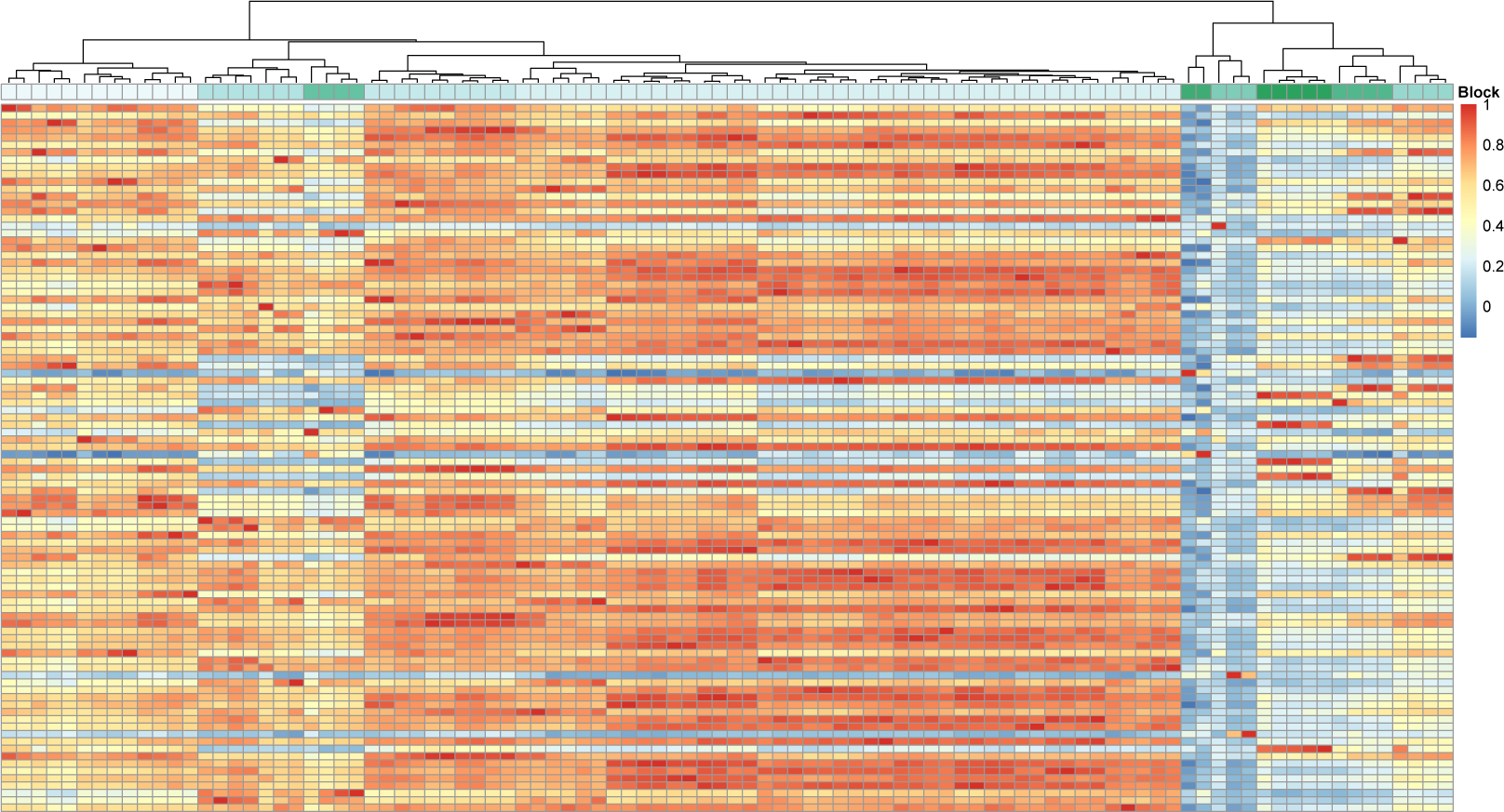
ADNI study: correlation map of the 96 regions of the brain. The 10 blocks obtained by hierarchical clustering for the PCEV-block approach can be identified using the dendrogram at the top of the figure.

## References

[1] Abdi H. Partial least square regression, projection on latent structure regression, PLS-Regression. Wiley Interdisciplinary Reviews: Computational Statistics, 2010.

[2] Härdle W and Simar L. Canonical correlation analysis. Applied Multivariate Statistical Analysis 2007;: 321–330.

[3] Friedman J. Regularized discriminant analysis. Journal of the American Statistical Association 1989; 84(405): 165–175.

[4] Ott J and Rabinowitz D. A principal-components approach based on heritability for combining phenotype information. Human heredity 1999; 49(2): 106–111.

[5] Wang Y, Fang Y and Man J. A ridge penalized principal-components approach based on heritability for high-dimensional data. Human Heredity 2007; 64: 182–191.

[6] Klei L, Luca D, Devlin B et al. Pleiotropy and principal components of heritability combine to increase power for association analysis. Genetic epidemiology 2008; 32(1): 9–19.

[7] Fang Y, Feng Y and Yuan M. Regularized principal components of heritability. Com-putational Statistics 2014; 29(3-4): 455–465.

[8] Nishisato S. Optimization and data structure: Seven faces of dual scaling. Annals of Operations Research 1995; 55(2): 345–359.

[9] Lin J, Zhu H, Knickmeyer R et al. Projection regression models for multivariate imaging phenotype. Genetic epidemiology 2012; 36(6): 631–641.

[10] Hoerl AE and Kennard RW. Ridge regression: Biased estimation for nonorthogonal problems. Technometrics 1970; 12(1): 55–67.

[11] Tibshirani R. Regression shrinkage and selection via the lasso. Journal of the Royal Statistical Society Series B (Methodological) 1996;: 267–288.

[12] Everitt B and Dunn G. Applied Multivariate Data Analysis. Edward Arnold. London, 1991.

[13] Rencher A and Christensen W. Methods of Multivariate Analysis. John Wiley and Sons, 2012.

[14] Johnstone IM. Multivariate analysis and jacobi ensembles: Largest eigenvalue, tracy-widom limits and rates of convergence. Annals of statistics 2008; 36(6): 2638.

[15] Cotterchio M, McKeown-Eyssen G, Sutherland H et al. Ontario familial colon cancer registry: methods and first-year response rates. Chronic Dis Can 2000; 21(2): 81–86.

[16] Zanke BW, Greenwood CM, Rangrej J et al. Genome-wide association scan identifies a colorectal cancer susceptibility locus on chromosome 8q24. Nature genetics 2007; 39(8): 989–994.

[17] Fortin JP, Labbe A, Lemire M et al. Functional normalization of 450k methylation array data improves replication in large cancer studies. Genome Biol 2014; 15(11): 503.

[18] Houseman EA, Accomando WP, Koestler DC et al. DNA methylation arrays as surrogate measures of cell mixture distribution. BMC bioinformatics 2012; 13(1): 86.

[19] Miceli-Richard C, Wang-Renault SF, Boudaoud S et al. Overlap between differentially methylated DNA regions in blood B lymphocytes and genetic at-risk loci in primary Sjäogren’s syndrome. Ann Rheum Dis 2015; 0(0): 1–8.

[20] Gao X, Starmer J, Martin ER et al. A multiple testing correction method for genetic association studies using correlated single nucleotide polymorphisms. Genetic epidemi-ology 2008; 32(4): 361.

[21] Breitling LP, Yang R, Korn B et al. Tobacco-smoking-related differential DNA methy-lation: 27K discovery and replication. The American Journal of Human Genetics 2011; 88(4): 450–457.

[22] Lee KW and Pausova Z. Cigarette smoking and DNA methylation. Frontiers in genetics 2013; 4.

[23] Formisano E, De Martino F and Valente G. Multivariate analysis of fMRI time series: classification and regression of brain responses using machine learning. Magnetic resonance imaging 2008; 26(7): 921–934.

[24] Kherif F, Poline JB, Flandin G et al. Multivariate model specification for fMRI data. Neuroimage 2002; 16(4): 1068–1083.

[25] Rowe DB and Hoffmann RG. Multivariate statistical analysis in fMRI. Engineering in Medicine and Biology Magazine, IEEE 2006; 25(2): 60–64.

[26] Teipel SJ, Born C, Ewers M et al. Multivariate deformation-based analysis of brain atrophy to predict Alzheimer’s disease in mild cognitive impairment. Neuroimage 2007; 38(1): 13–24.

[27] Lindenmayer JP, Bernstein-Hyman R, Grochowski S et al. Psychopathology of schizophrenia: initial validation of a 5-factor model. Psychopathology 1995; 28(1): 22–31.

[28] Livshits G, Roset A, Yakovenko K et al. Genetics of human body size and shape: body proportions and indices. Annals of human biology 2002; 29(3): 271–289.

[29] Arya R, Blangero J, Williams K et al. Factors of insulin resistance syndrome-related phenotypes are linked to genetic locations on chromosomes 6 and 7 in nondiabetic mexican-americans. Diabetes 2002; 51(3): 841–847.

[30] Johnstone IM. Approximate null distribution of the largest root in multivariate analysis. The annals of applied statistics 2009; 3(4): 1616.

[31] Tracy CA and Widom H. On orthogonal and symplectic matrix ensembles. Communi-cations in Mathematical Physics 1996; 177(3): 727–754.

[32] Barker M and Rayens W. Partial least squares for discrimination. Journal of chemo-metrics 2003; 17(3): 166–173.

